# Assessing graph-based read mappers against a novel baseline approach highlights strengths and weaknesses of the current generation of methods

**DOI:** 10.1101/538066

**Authors:** Ivar Grytten, Knut D. Rand, Alexander J. Nederbragt, Geir K. Sandve

**Affiliations:** Department of informatics, University of Oslo, Oslo, Norway; Department of Mathematics, University of Oslo, Oslo, Norway; Department of Biosciences, University of Oslo, Oslo, Norway

## Abstract

Graph-based reference genomes have become popular as they allow read mapping and follow-up analyses in settings where the exact haplotypes underlying a high-throughput sequencing experiment are not precisely known. Two recent papers show that mapping to graph-based reference genomes can improve accuracy as compared to methods using linear references. Both of these methods index the sequences for most paths up to a certain length in the graph in order to enable direct mapping of reads containing common variants. However, the combinatorial explosion of possible paths through nearby variants also leads to a huge search space and an increased chance of false positive alignments to highly variable regions.

We here assess three prominent graph-based read mappers against a novel hybrid baseline approach that combines an initial path determination with a tuned linear read mapping method. We show, using a previously proposed benchmark, that this simple approach is able to improve accuracy of read-mapping to graph-based reference genomes.

Our method is implemented in a tool Two-step Graph Mapper, which is available at https://github.com/uio-bmi/two_step_graph_mapper along with data and scripts for reproducing the experiments.

## Background

Graph-based reference genomes are increasingly used as references in next-generation sequencing experiments, such as in variant calling [1, 2] and peak calling [4]. A key step in all such analysis pipelines is the alignment of raw sequencing reads to the reference. Recently, two tools for mapping reads to graph-based reference genomes have been proposed – *vg* [2] and a tool created by *Seven Bridges* [5] (from here on we refer to this tool as Seven Bridges). Both show improved mapping accuracy compared to the linear reference-based method BWA-MEM [6]. While *vg* indexes all paths up to a certain length in the graph – a tedious process that takes more than a day for a human whole-genome graph – Seven Bridges uses a faster approach in which only short kmers (21 base pair sequences at 7 base pairs intervals) are indexed. This enables indexation of a human whole-genome graph in only minutes. Both methods ignore the most complex regions in the graph in order to reduce the number of kmers to index. A third method for mapping reads to graph-based references is Hisat 2, which uses a Hierarchical Graph FM index [3].

There currently exists no comparison of the mapping accuracy of *vg*, Seven Bridges and Hisat 2. Furthermore, there exists no study on how these tools perform compared to linear mapping approaches tuned for accuracy and not speed, or to simpler schemes for graph-based read mapping. We here present a novel hybrid graph-mapping approach and use this as a baseline to highlight strengths and potential for improvement for the current generation of graph-based mapping approaches. We compare *vg,* Seven Bridges and Hisat 2 to a tuned linear mapping approach, and to our novel two-step approach, and show that graph-based read mapping can be improved by separating the problem into rough path estimation and subsequent mapping of each individual read to this estimated path.

## Results

In the following, we assess graph-mappers by looking at *vg*, Seven Bridges and Hisat 2. All assessments are done by following the approach that *vg* and Seven Bridges used for evaluating their tools [5]. We simulate 10 million single-end reads from chromosome 20 of an Ashkenazi Jewish male NA24385, sequenced by the Genome in a Bottle Consortium [8] (see Methods). We simulate uniformly across the genome, and some reads will naturally be simulated from segments containing non-reference alleles (about 10.6% of the reads). We refer to these as *reads with variants.* Reads that are simulated from segments identical to the linear reference genome (hg19) will be referred to as *reads without variants.* Mapping accuracies are compared using ROC-plots parameterized by the mapping quality of all the simulated reads. Scripts and data generating the figures in this section are provided at https://github.com/uio-bmi/two_step_graph_mapper.

### *vg* outperforms Seven Bridges and Hisat 2 on previously proposed benchmarks

In Figure 1, we compare the mapping accuracy of *vg*, Seven Bridges and Hisat 2 on the simulated reads, using two different ways of simulating reads – using *vg sim* with 1% substitution rate and 0.2% indel rate as used by *vg* in their evaluation [2] and by using the tool Mitty with an error model giving on average 0.26% substitution rate, as used by Seven Bridges in their evaluation [5]. *vg* performs better than both Seven Bridges and Hisat 2 on both error rates. From here on, we thus focus on *vg* when discussing capabilities and limitations of the current generation of graph-based mapping approaches, and use simulated reads with 1% substitution rate and 0.2% indel rate (as used by *vg* in their evaluation).

**Figure 1.**
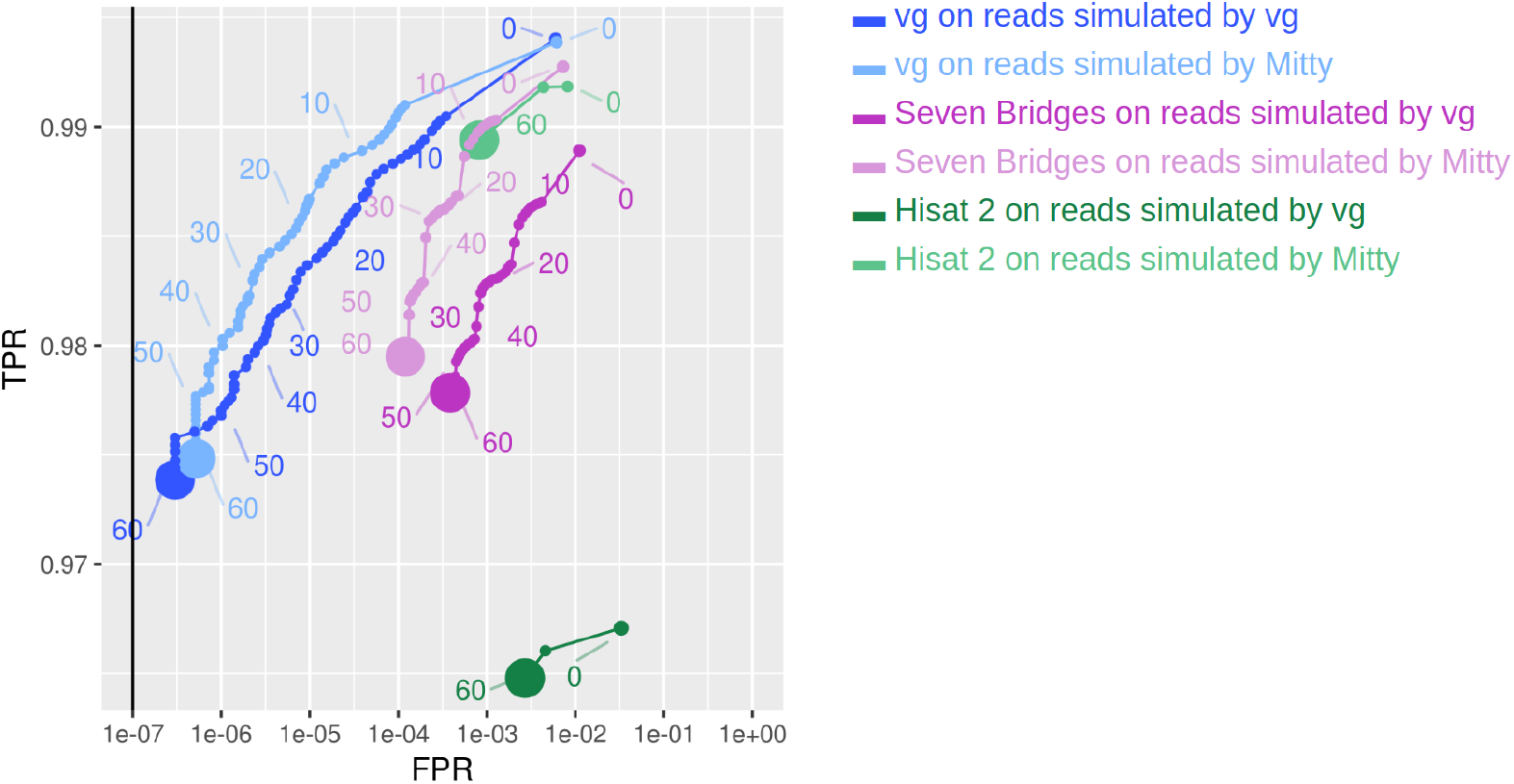
Comparison of mapping accuracy on reads mapped by *vg* and Seven Bridges by ROC-plots parameterized by the mapping quality of reads simulated two ways: Using *vg sim* with base pair substitution rate 1% and indel rate 0.2% (what *vg* used when evaluating their methods) and by using Mitty to simulate reads using the error model “hiseq-2500-v1-pcr-free”, which is what Seven Bridges used in their evaluation (average substitution rate of 0.26%).

### Part of the performance difference between graph-based and linear methods can be attributed to method tuning

As shown in Figure 2, *vg* performs better than BWA-MEM when BWA-MEM is run with default parameters. However, BWA-MEM is by default tuned for speed and not for maximum accuracy. By tuning BWA-MEM and adjusting the MAPQ scores by also running Minimap 2 (see Methods), BWA-MEM goes from performing worse than *vg* on all reads to be performing about as well as *vg*. From here on, we use this tuned version of BWA-MEM, referred to as *linear mapper,* when comparing graph-based and linear mapping approaches.

**Figure 2.**
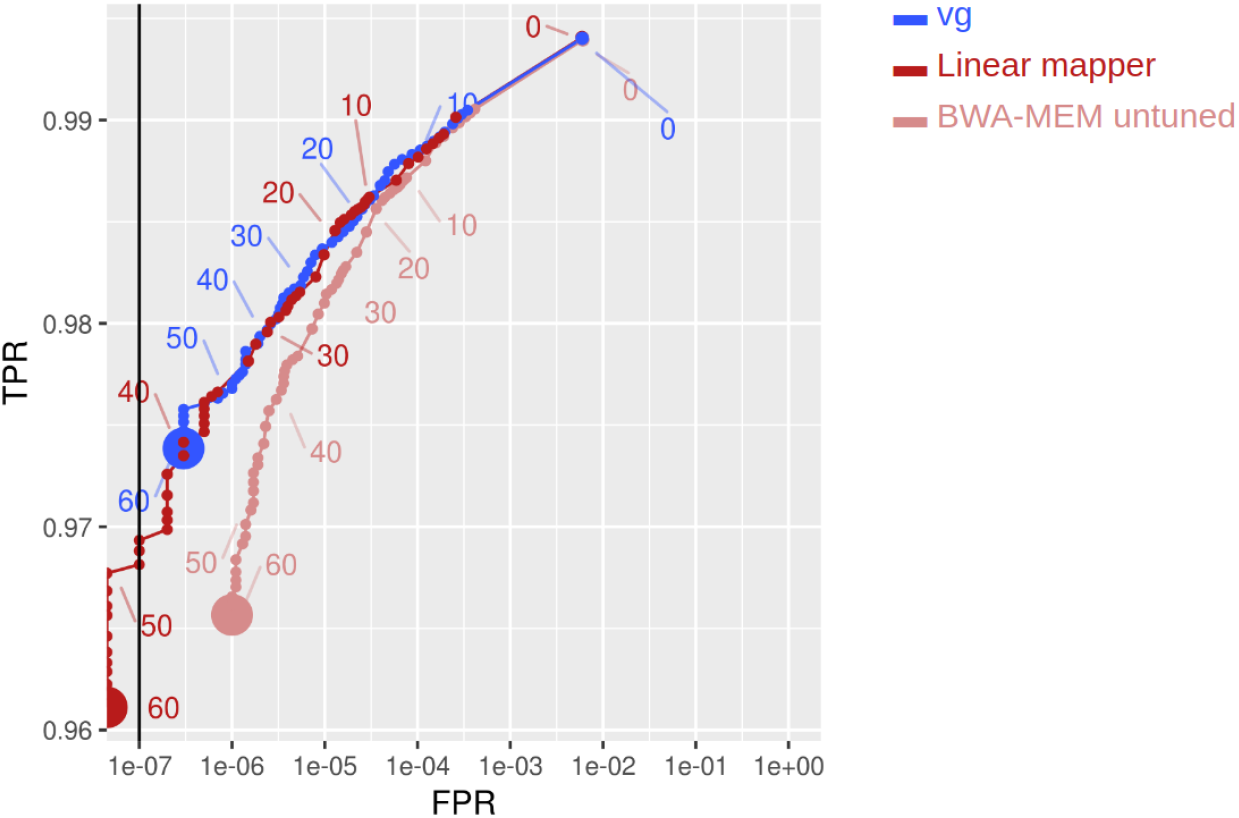
Comparison of the mapping accuracies of “Linear mapper” (BWA-MEM with tuned parameters using Minimap 2 to adjust MAPQ scores), *vg* and untuned BWA-MEM (running with default parameters).

### Graph-based mapping results in higher accuracy on reads with variants, but lower accuracy on reads without variants

As seen in Figure 3, *vg* achieves markedly higher accuracy on reads with variants than the linear mapper (tuned BWA-MEM). However, as also noted in [2], the mapping accuracy of *vg* is lower than the linear mapper on reads that do not contain variants. As a result of this, *vg* ends up not performing better than the linear mapper when assessed on the full set of reads.

**Figure 3.**
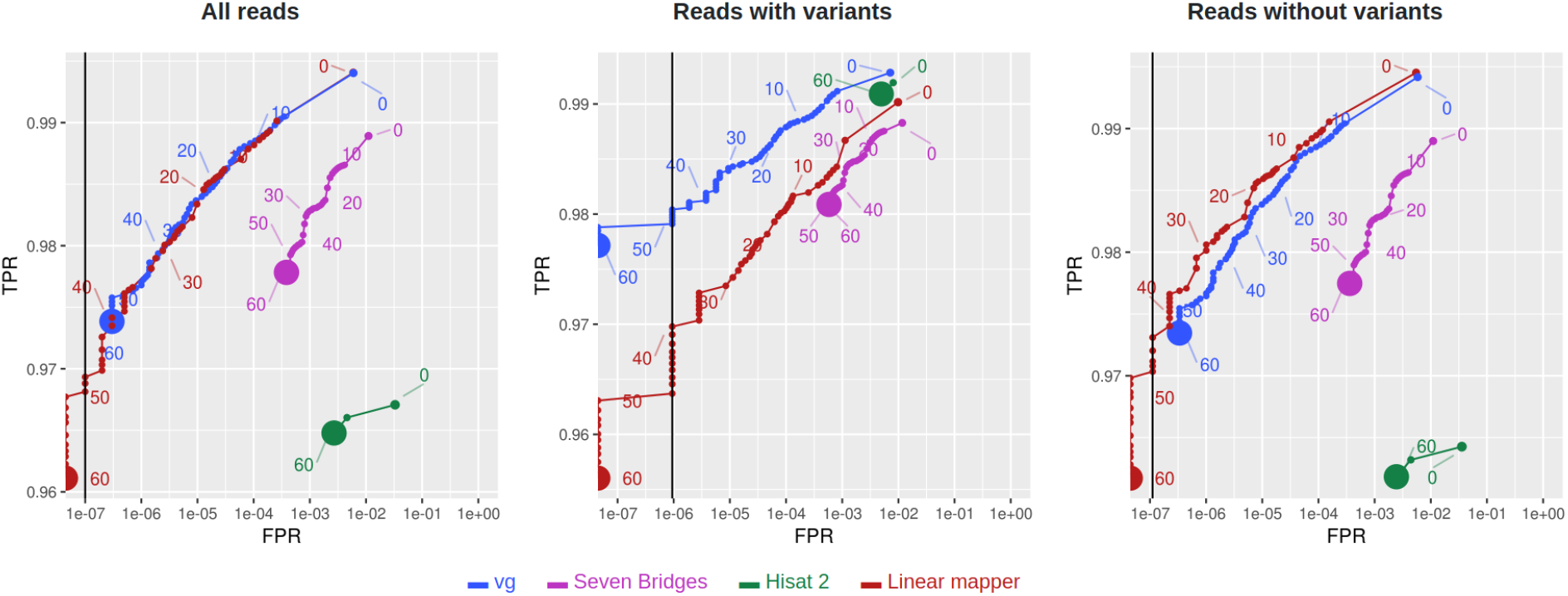
Comparison of mapping accuracies of *vg,* Seven Bridges and linear mapping (tuned BWA-MEM).

### Re-aligning the reads to an estimated linear path through the graph improves accuracy

We find that using the initial graph alignments to predict a linear path through the graph, and then re-aligning all the reads to this linear path using the linear mapper increases mapping accuracy. In Figure 4, we show the results of this approach. As seen in the figure, this two-step approach performs almost as well as *vg* on reads containing variants and clearly better than *vg* on reads not containing variants, resulting in better overall performance on all reads.

**Figure 4.**
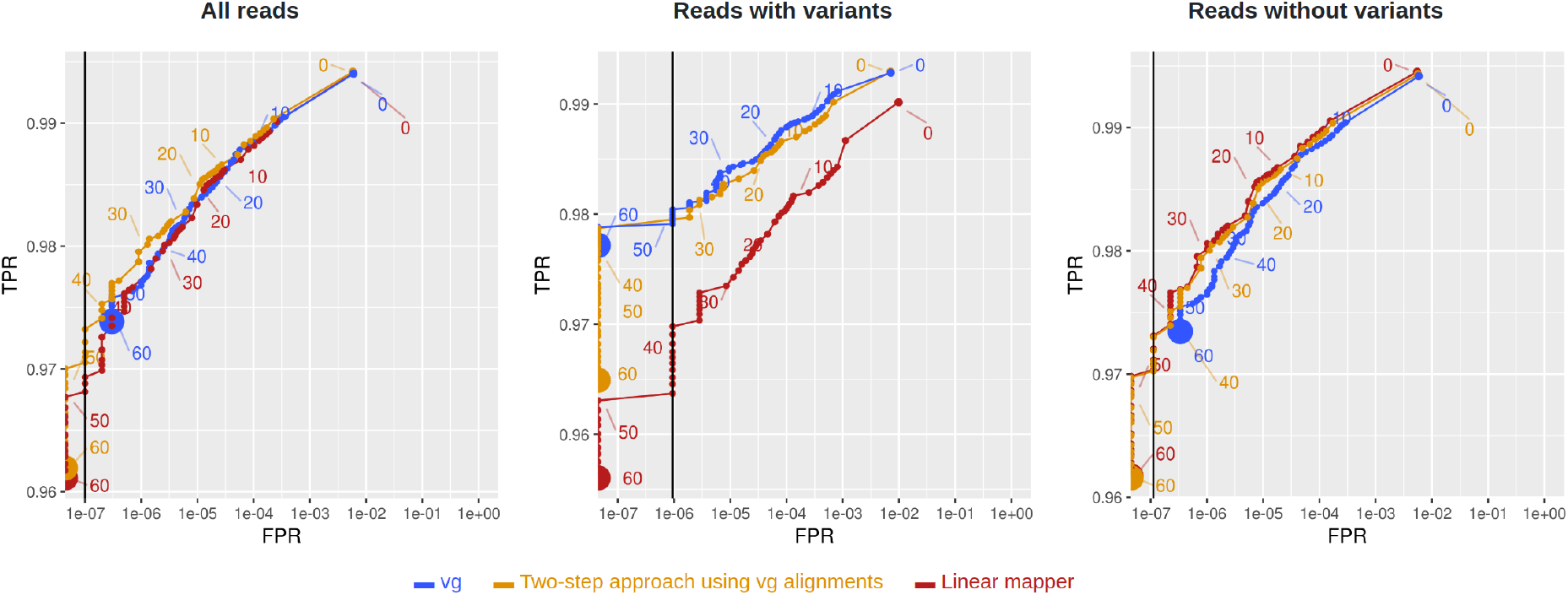
Comparison of *vg,* the linear mapper and a two-step approach using *vg* alignments to initially predict a path through the graph and then re-aligning the reads to this path using the linear mapper.

### A novel two-step approach using an initial rough path estimation is sufficient to improve mapping accuracy

The results from the previous section indicate that the *vg* mapping accuracy can be improved by predicting a path and re-aligning all the reads to this path using the linear mapper. We argue that this idea works as long as we are able to predict an approximate path in the first step. We suggest that the path-prediction in itself can be achieved by initial rough graph-mapping, and as an example, we use an initial rough graph-mapping method where all the reads first are aligned to the linear reference genome and then subsequently re-aligned locally to the graph using using Graph SIMD Smith-Waterman (GSSW) [7]. A proof-of-concept implementations of this method is provided in the Python package Rough Graph Mapper (https://github.com/uio-bmi/rough_graph_mapper).

As seen in Figure 5, the use of this method in the first step of the two-step approach leads to better mapping accuracy than *vg* for non-variant reads, and almost as good accuracy as *vg* on variant-reads. Since this is possible with an initial rough mapping that does not rely on a graph index (like the one used by vg) we argue that this two-step approach is a promising direction for computationally efficient graph-based read mapping. Our two-step approach is implemented in a tool Two-step Graph Mapper, which is available at https://github.com/uio-bmi/two_step_graph_mapper.

**Figure 5.**
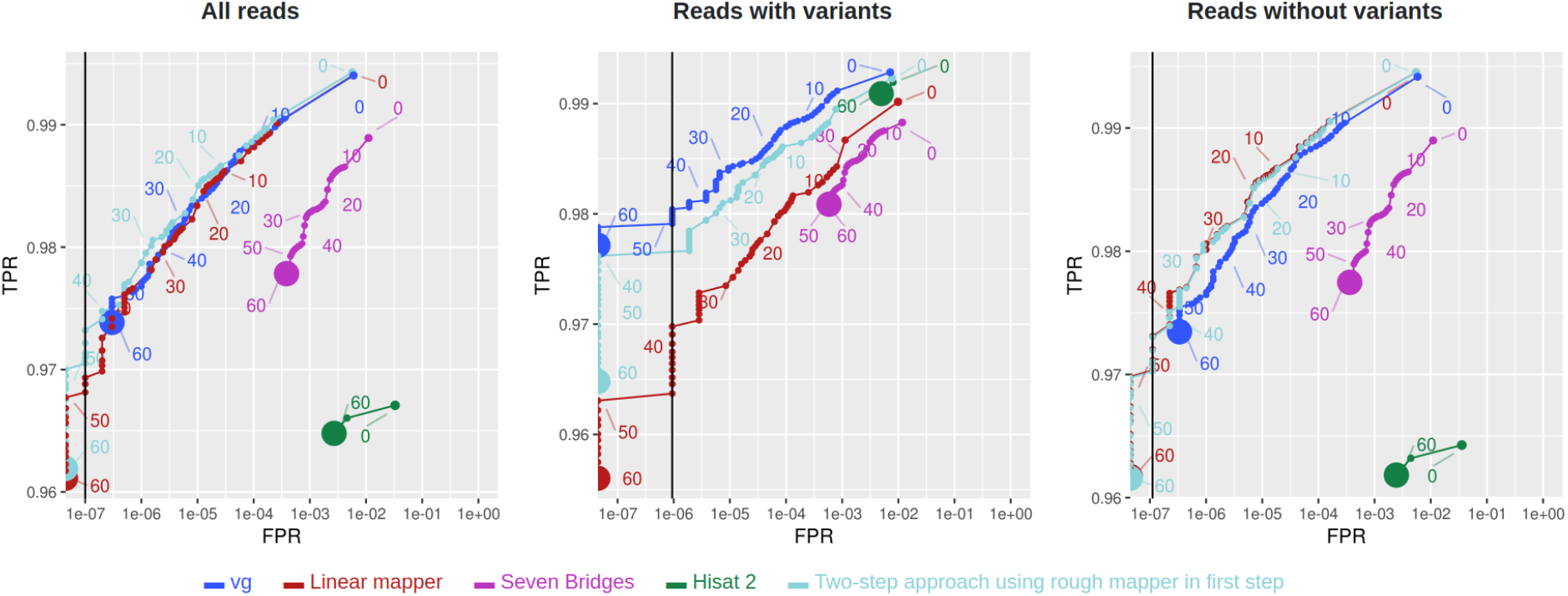
Comparison of mapping accuracies of the two-step approach using two methods for rough initial alignment, the linear mapper, *vg* and Seven Bridges.

## Discussion

We observe higher accuracy for *vg* than Seven Bridges and Hisat 2 in our comparisons. We feel these comparisons are an unfortunate omission in the Seven Bridges paper. These three methods all perform worse than linear mapping on reads not containing variants, and a tuned version of BWA-MEM achieves about the same accuracy as *vg* on the full set of reads. We are unsure why Hisat 2 performs worse than vg, but to our knowledge, Hisat 2 is primarily used for RNA and not DNA sequencing reads. We hypothesise that Seven Bridges performs worse than vg because it is using a much simpler index, containing only a subset of all kmers in the graph. We further show that a two-step approach of predicting a path through the graph and mapping to this path using the linear mapper results in higher accuracy, even when using a rough graph-mapper for the initial prediction of the path. Our two-step approach achieves about the same accuracy as *vg* on reads containing variants and higher accuracy than *vg* on reads not containing variants (which contribute to about 90% of the simulated reads). We believe this is because the use of a predicted path limits the search space dramatically in the final mapping.

Read alignment serves as an intermediate step for several distinct investigations. The aligned reads may be used as input for variant callers in order to determine genotypes or somatic mutations, for peak callers to determine locations of epigenetic modifications or protein binding to DNA, and for transcriptome analysis methods to quantify differential gene expression or alternative splicing. The consequences of different categories of mis-mapped reads (e.g. reads originating from genomic regions of high or low variation) may vary between these settings. We have in this work restricted our focus to assessing the accuracy of read mapping. It would be interesting to explore how the mis-mapping profiles of the different approaches affect the following analysis step for each such setting.

We have shown one implementation of how reads can be mapped in the first step of the two-step approach. This method maps each read to the linear reference genome first and then locally re-aligns each read to the graph. A variant of this method that probably would give better results would be to have the linear mapper report the *n* best hits for each read, locally align each of those to the graph, and pick the alignment with highest graph alignment score. As future work, we also believe it could be interesting to use other graph-based mapping methods that sacrifice accuracy for speed in the first step in the two-step mapping approach. An idea for such a method could be a graph-generalization of minimizer-based mapping methods such as Minimap [10].

The method we use for initial rough path prediction is fairly simple and naive, but illustrates the point. As future work, it would be interesting to implement more sophisticated path prediction algorithms, e.g. including haplotype information or correlations between variants in the graph. We note that our two-step approach only performs well when there are sufficient reads for predicting the path (i.e. high enough coverage). In the simulations, we deliberately only simulated reads from one chromosome in order to get high enough coverage (20x) without needing hundred of millions of reads. The coverage used (20x) is lower than than the typical coverage for most modern next generation sequencing experiments, such as whole genome sequencing, RNA-seq and the coverage within peaks in a ChIP-seq experiment. With lower coverage, we expect the accuracy to drop down to that of a linear sequence aligner, since our path prediction algorithm defaults to the linear reference genome path when there is not enough reads covering a variant. With higher coverage, we expect the accuracy to increase. Our current implementation only predicts one path through the graph, but in reality, reads coming from a diploid individual will follow two paths. It should be trivial to instead estimate two paths in the first step of our two-step approach, and align reads to both paths in the final step.

For linear reference genomes, the sole objective of mapping is to align reads back to the genomic locations they originate from. In contrast, mapping against graph-based reference genomes can serve a dual purpose: estimating the underlying haplotypes (two paths through the graph) and correctly placing each read along these haplotype paths. The driving idea of our two-step approach is to separate these as two separate algorithmic problems. This allows a rough mapping approach to be used initially for estimating the haplotype and thus limit the search space for a subsequent step of placing reads along this path using any linear mapper. It is important to note that although the path-estimation in the first step of the two-step approach implicitly estimates variants present in the graph, the intention of this step is not to do variant calling – instead variant calling can be performed as a follow-up step based on the aligned reads.

## Conclusions

We have here proposed a hybrid baseline approach for graph-based read mapping that combines an initial path determination with a tuned linear read mapping method. By comparing three prominent graph-based read mappers to this novel baseline, we find that part of the accuracy gains observed in recent comparisons of graph-based and linear mappers can be attributed to method tuning. Nonetheless, when focusing on reads containing variants (as compared to the linear reference genome), we observe markedly improved accuracy of the graph-based mapper *vg* as compared to mapping to a linear reference using a tuned version of BWA-MEM. Two other graph-based mappers, Seven Bridges and Hisat 2, attain markedly lower mapping accuracy than *vg* in our benchmarks, and do not improve on the linear mapper even on the regions containing variants. By employing *vg* for initial path determination in our proposed two-step approach, we improve on the performance of *vg* used in isolation. Furthermore, even when using a quick, rough mapper for the initial step, our two-step approach performs comparably to the use of *vg* in isolation. In addition to serving as a baseline for highlighting characteristics of the current generation of graph-based read mappers, we thus believe that our two-step approach represents a promising alternative direction for computationally efficient graph-based read mapping.

## Methods

### Assessment of mapping methods

We compared *vg*, Seven Bridges and Hisat 2, which to our knowledge are the only published tools for mapping reads to graph-based reference genomes. We considered to include running BWA-MEM on an index created by CHOP [9], which is a tool for indexing paths through known haplotypes in the graph, but we were unable to build the CHOP index for a whole human genome human graph in reasonable run-time. We also considered to include PanVC [11] in the comparison, but also this method does not seem to scale to a whole human genome, for which it takes weeks to run on [11]. When using vg to simulate reads, we used base pair substitution rate 0.01 and indel rate 0.02. When using Mitty, we used the built-in error model hiseq-2500-v1-pcr-free. All evaluations were run with vg version 1.12.1, BWA-MEM version 0.7.17-r1194-dirty, Minimap 2 version 2.13, Seven Bridges graph aligner bpa-0.9.1.1-3, Hisat 2.1.0 and Mitty version 2.28.3. We ran vg and Seven Bridges with default parameters and Hisat 2 with the –no-spliced-alignment option. When running BWA-MEM, we tuned it and used Minimap in order to to adjust the MAPQ scores returned by BWA-MEM (see the next section for details).

The benchmark we ran is similar to the one performed by vg in [2] and the one by Seven Bridges in [5]. In all analyses, we used a graph created from variants from the 1000 Genomes Project having allele frequency > 1% (about 14 million variants), since this graph gave the best results for vg and seems to be similar to the graph that Seven Bridges used in their evaluation, which contained 15.8 million variants (mostly from the 1000 genomes project).

All reads were simulated from chromosome 20 only, in order to get high coverage without having to simulate very many reads. We only simulated single-end reads. ROC-plots were generated by running the same scripts used by vg to create ROC-plots in [2]. We have created a Docker image with all scripts and exact software with dependencies necessary for re-running the benchmarks. This image along with instructions on how to re-run the analysis are available from the readme page at https://github.com/uio-bmi/two_step_graph_mapper.

### Tuning BWA-MEM performance

We observe worse performance by BWA-MEM with default parameters compared to vg, even on reads not containing variants. This has also been noted in [2] where vg running on a linear reference genome is shown to outperform BWA-MEM running on the same reference. From our experience, a cause of this is that BWA-MEM is tuned for speed (not only performance), mainly by not trying to align shorter “chains” by default. This makes BWA-MEM miss a lot of suboptimal alignments (when they exist), which in turn makes it overestimate the MAPQ score. Changing the -D parameter of BWA-MEM to a low number partly solves this, by telling BWA-MEM to also try to align shorter chains. However, after such tuning, there are still cases where BWA-MEM fails to find suboptimal alignments. This typically happens when all the longest chains cover a sequencing error in the read. We did not find a way of tuning BWA-MEM to consider shorter chains in such cases, but we found that Minimap 2 (even though it generally performs worse than BWA-MEM on short reads) in most cases was able to find all suboptimal alignments and use that to correctly assign low MAPQ scores when reads were multimapping. Thus, we chose to also run Minimap 2 on the same reads and for every read, simply selecting the MAPQ score chosen by Minimap 2 when Minimap 2 assigned a lower MAPQ score than BWA-MEM.

We did not tune vg in order to try to improve its performance, since it seems that vg with default parameters is already tuned to perform well (we did not see any cases of vg missing alignments that BWA-MEM found).

### Two-step graph mapping

#### Step 1: Estimating a path through the graph

In order to predict a path through the graph, some initial graph-alignments are needed. We have proposed a simple way for performing this initial mapping, which is explained more in detail in the next section. In this first step, we use the rough graph alignments to predict a path through the graph. We do this by greedily traversing the graph by always following the edges with most reads aligned to them (edges going to linear reference nodes are given a bonus, default is 1). We then extract the sequence of this path, and use BWA (bwa index with default parameters) to index it.

#### Step 2: Aligning the reads

In this step, we simply align all the reads to the indexed path using the tuned version of BWA-MEM. After the reads have been aligned to this predicted path, their coordinates will be relative to this path, thus differing from the linear reference genome (e.g. hg19) coordinates. We have implemented a simple method for translating the coordinates back to linear reference genome coordinates (similar to vg annotate) that simply moves the alignments to the graph and finds the closest position on the linear reference path going through the graph.

#### Rough mapping of reads to the graph

First, we map all the reads to the linear reference genome using the linear mapper (tuned BWA-MEM). We then move these reads to their corresponding position in the graph, which is possible since the graph contains the linear reference genome. We then re-align each read locally to the graph using the Graph SIMD Smith-Waterman package [7].

